# A novel barcoded nanopore sequencing workflow of high-quality, full-length bacterial 16S amplicons for taxonomic annotation of bacterial isolates and complex microbial communities

**DOI:** 10.1101/2024.04.11.588846

**Authors:** Julian Dommann, Jakob Kerbl-Knapp, Diana Albertos Torres, Adrian Egli, Jennifer Keiser, Pierre H. H. Schneeberger

## Abstract

**Introduction:** Due to recent improvements, Nanopore sequencing has become a promising method for experiments relying on amplicon sequencing. We describe a flexible workflow to generate and annotate high-quality, full-length 16S rDNA amplicons. We evaluated it for two applications, namely, i) identification of bacterial isolates and ii) species-level profiling of microbial communities.

**Methods:** Bacterial isolate identification by sequencing was tested on 47 isolates and compared to MALDI-TOF MS. 97 isolates were additionally sequenced to assess the resolution of phylogenetic classification. Species-level community profiling was tested with two full-length 16S primer pairs (A and B) with custom barcodes and compared to results obtained with Illumina sequencing using 27 stool samples. Finally, a Nextflow pipeline was developed to produce high-quality reads and taxonomically annotate them.

**Results:** We found high agreement between our workflow and MALDI-TOF data for isolate identification (PPV = 0.90, Cramér’s V = 0.857 and, Theil’s U = 0.316). For species-level community profiling, we found strong correlations (r_s_ > 0.6) of alpha diversity indices between the two primer sets and Illumina sequencing. At the community level, we found significant but small differences when comparing sequencing techniques. Finally, we found moderate to strong correlation when comparing relative abundances of individual species (average r_s_ = 0.6 and 0.533, for primers A and B).

**Discussion:** The proposed workflow enabled accurate identification of single bacterial isolates, making it a worthwhile alternative to MALDI-TOF. While shortcomings have been identified, it enabled reliable identification of prominent features in microbial communities at a fraction of the cost of Illumina sequencing.

**Importance:** A quick, robust, simple, and cost-effective method to identify bacterial isolates and communities in each sample is indispensable in the fields of microbiology and infection biology. Recent technological advances in Oxford Nanopore Technologies sequencing make this technique an attractive option considering the adaptability, portability, and cost-effectiveness of the platform. Here, we validated a flexible workflow to identify bacterial isolates and characterize bacterial communities using the Oxford Nanopore Technologies sequencing platform combined with the most recent v14 chemistry kits. For bacterial isolates, we compared our nanopore-based approach to MALDI-TOF MS-based identification. For species-level profiling of complex bacterial communities we compared our nanopore-based approach to Illumina shotgun sequencing. For reproducibility purposes, we wrapped the code used to process the sequencing data into a ready-to-use and self-contained Nextflow pipeline.

## Introduction

Identifying the interaction between bacterial communities and their environment has shed light on the importance of bacterial communities in both health and disease. The potential to utilize microbes as research subjects to understand, identify and treat maladies helped drive innovative techniques such as Oxford Nanopore Technology (ONT) sequencing, to allow high-throughput identification of bacteria. Currently, among the different methods to analyze DNA, ONT sequencing has emerged as a novel approach. Although initial drawbacks in accuracy (few correctly aligned bases between generated sequences and reference genomes) limited the robustness and applicability of this method, continuous advances in kit-chemistry, flow cell design, and available bioinformatics tools have drastically increased the usability of the ONT sequencing platform (1–4). Improvements in base-calling yielded high-quality reads, whereas automatable data processing allowed simultaneous processing of large numbers of samples. With the recent v14 chemistry kits, an accuracy of 99.9% (Phred score of 30) is achievable, turning it into a robust, affordable, and particularly versatile option in the field.

Recently, Urban et al. demonstrated the feasibility of utilizing the ONT platform for the investigation of large numbers of bacterial samples. They employed custom primers to add barcodes during 16S rDNA PCR amplification, significantly reducing the cost per sample to approximately 20 GBP (5). Building upon this approach, it is desirable to develop a workflow that is easily implementable, cost-effective, and adaptable to diverse microbiological samples with potential high-throughput applications.

Hereby, we firstly tested and optimized a step-by-step workflow for either a single (1BC) or double barcoding (2BC) approach using ONT sequencing (v14). Secondly, we developed a flexible data processing pipeline, involving either simplex (S1BC or S2BC) or duplex (D1BC or D2BC) base-calling, depending on the application’s needs (e.g. sequencing accuracy versus sequencing depth). The code is available as a Nextflow pipeline based on validated bioinformatics tools enabling filtering, quality control, trimming, demultiplexing, and taxonomic annotation of the sequenced 16S rDNA amplicons. We therefore aim to fill the gap for an easily implementable, cost-effective, and adaptable workflow and analytical pipeline to study diverse microbiological specimens using ONT sequencing.

## Materials and methods

### Samples

We initially sequenced 47 bacterial isolates, for which identification results via MALDI-TOF MS mass spectrometry (MS) were available. We sequenced another 97 bacterial isolates, to further explore the potential of our sequencing approach regarding phylogenetic resolution. In total we therefore sequenced 144 bacterial isolates, spanning across 41 species and 20 genera. While not all isolates are of medical relevance, we enhanced taxonomic diversity by including both phylogenetically distant and close species, both aerobic and anaerobic species and isolates originating from different sample types (for instance from blood or stool). The 144 bacterial isolates tested in this study are listed in **Supplementary Table 1**. Bacterial 20% glycerol stocks were stored at -80℃. To assess performance of species-level microbiome characterization, we sequenced DNA extracts of 27 human stool samples gathered from healthy individuals. Stool samples were gathered as part of a multi-country randomized controlled trial (NCT03527732; https://clinicaltrials.gov/ct2/show/NCT03527732) with the goal of evaluating the efficacy and safety of antiparasitic medications against *T. trichiura* (6). Individuals interested in participating in the trial were invited to complete the process of informed consent; thereafter, individuals were assessed for study eligibility during screening procedures. Prior to the start of collection, participants were informed of the aim of the daily sample collection, in addition to the consenting and information sessions conducted for the trial. For each participant, a small aliquot of stool (∼1g) was transferred to a sterile 2 mL cryotube and immediately frozen at -20°C. Upon completion of the trial, these aliquots were shipped to Swiss TPH (Allschwil, Switzerland) on dry ice and kept at -80°C until DNA extraction. Only fully anonymized samples and isolates were used throughout the study and no participant-specific information was available.

### Bacterial Cultivation

Growth media were prepared using Milli-Q water and subsequently autoclaved at 121°C. Brain Heart Infusion broth (BHI) + 5% Yeast or modified Gifu anaerobic medium (mGAM) were used exclusively to cultivate all bacterial isolates. To cultivate an isolate, 10μl of thawed glycerol stock was used to start a culture in 10ml growth medium. To cultivate under anaerobic conditions, a vinyl anaerobic chamber (Coy Laboratory Products, Michigan, United States) with a gas mix composed of 85% N_2_, 10% CO_2_ and 5% H_2_ was used. Prior to working, the anaerobic chamber was cleaned using a 1:750 dilution of benzalkonium chloride (distribution from C_8_H_17_ to C_16_H_33_) in purified water to avoid cross-contamination. Aerobic isolates were handled in a safety cabinet (SKAN Berner, Elmshorn, Germany). All inoculated isolates were grown at 37°C. Growth of isolates was inspected daily, turbid cultures were pelleted at 3.000 rcf, supernatant removed up to 1ml, and stored at -20°C.

### DNA extraction

Bacterial pellets were thawed at room temperature and DNA was extracted using a commercially available extraction kit (DNeasy® PowerSoil ® Pro Kit, QIAGEN, Germany) according to the manufacturer’s protocol. Deviating from the protocol, 200 µL resuspended bacterial pellet was used as input, and the DNA was ultimately eluted in 60 µL C6 elution buffer to maximize yield while keeping the DNA concentration high. For the microbiome samples, ∼100 mg of the stool sample was processed with the same kit using the same steps. DNA concentration and purity of all samples was measured using a Qubit 4 Fluorometer and the dsDNA BR Assay kit (both Invitrogen, USA)

### Barcoded 16S rDNA amplification

To acquire barcoded amplicons at high concentration, full-length 16S rDNAs of bacterial isolates or stool samples were amplified in a 96-well plate using barcoded 16S primers. The extracted DNA was first diluted 1:10 in nuclease-free water (Ultrapure™ Distilled water, Invitrogen, USA). The underlying design of our 16S primer sequences was published by Urban *et al*. 2021 (5). Our adapted primers (A) contain an added barcode sequence (see **Supplementary Table 2**), that was derived from Illumina 12bp barcode sequences published by Caporaso *et al.* 2012 (7). We also tested full-length 16S primers, recently published by Matsuo et al. (8) utilizing degenerate bases to resolve PCR bias and taxonomic underrepresentation of *Bifidobacterium* spp. (B) (**Supplementary Table 3**). A and B primers were ordered from Microsynth (Balgach, Switzerland). For amplicon generation, a commercially available PCR Master mix was used (LongAmp® Hot Start *Taq* 2X Master Mix, New England BioLabs, USA). For each run, eight forward and reverse primer pairs were used yielding unique barcodes for each sample per plate column. Per-well reagents are presented in **Supplementary Table 4**. The reaction was performed on an Eppendorf Mastercycler Nexus Gradient Thermal Cycler (Eppendorf, Germany) using a ramp rate of 1.5℃/s and cycling conditions as described in **Supplementary Table 5**. DNA concentration of samples post-PCR was measured with the dsDNA BR Assay kit on a Qubit 4 Fluorometer (both Invitrogen, USA) according to the manufacturer’s protocol, using 2µL of PCR reaction as input. Furthermore, presence of PCR products and size were checked via gel electrophoresis in 1% agarose gels (100V, 25min). Sample plates were either stored at 4°C overnight or at -20°C for long-term storage.

### ONT library preparation and sequencing

Library preparation was done according to the protocol for the Native Barcoding Kit 24 v14 (SQK-NBD114.24, Oxford Nanopore Technologies, UK) to ensure higher sample input (opposed to the SQK-NBD114.96 protocol). We applied the same library preparation protocol for both, bacterial isolates as well as the stool samples. Steps deviating from the protocol aimed at normalizing the sample input, purifying short fragments, and allowing multiplexing. They were as follows: (1) Ahead of the end-repair step, samples were pooled column-wise from the 96-well plate. Thus, for 96 samples, 12 different outer barcodes needed to be used, one for every 8 samples. A total of 200ng - corresponding to 200 fmol of the expected 1.6kb fragments - was used as input for the end-repair steps. Equimolar amounts of each of the eight samples per column were used as input and filled up to 11.5µL with nuclease-free water and processed as one sample. (2) After the end repair, samples were measured using the 1X dsDNA HS Assay kit (Invitrogen, USA) on a Qubit 4 fluorometer. Equimolar masses of the individual pooled reactions were used as input for the native barcode ligation, which should lead to equal read numbers per barcode. Dilutions were made with nuclease-free water. (3) The adapter ligation and clean-up were done according to protocol, the short-fragment buffer was used to preserve the 16S rDNA fragments. (4) The protocol was modified for usage with a Flongle Flow Cell (Flongle Flow Cell, R10 Version, FLO-FLG114, Oxford Nanopore Technologies, UK) as follows: for the priming of the Flongle Flow Cell, 3 µL Flow Cell Tether was added to 117 µL Flow Cell Flush. The DNA concentration of the finished library was measured using the Qubit 4 fluorometer (1X dsDNA HS Assay kit) and diluted to a concentration of 2-3 ng/µL using nuclease-free water. For the loading of the library, 15 µL Sequencing buffer, 10 µL Loading Beads and 5 µL of the final library were mixed and fully loaded onto a Flongle Flow Cell on a MinION Mk1C (Oxford Nanopore Technologies, UK). An overview of the sequencing run settings is described in **Supplementary Table 6**. For the microbiome samples, each library was sequenced in triplicate on independent flow cells to mitigate sequencing bias and depth variations arising from flow-cell-specific differences in total sequencing output.

### Data processing and taxonomic annotation of Nanopore data

Read processing for bacterial isolate sequencing was done as follows: The POD5 files generated for each sequencing run were demultiplexed based on the native ONT barcode in real-time utilizing the MinION’s built-in Guppy (v6.5.7). The demultiplexed POD5 files were base-called using Dorado (v. 0.2.4; base-caller model “dna_r10.4.1_e8.2_400bps_hac@v4.1.0”; either simplex or duplex). Base-called reads served as input for a custom Nextflow (9) pipeline performing the following steps: First, reads are filtered based on quality and length using NanoFilt (10) (v. 2.8.0; 1300-1800bp, quality score > 9). Next, a MultiQC (11) (v. 1.11) report is generated to estimate the robustness and usability of every run via the mean quality score (Q-score), average read length and average read numbers. Subsequently, a combination of seqkit (v.2.6.1) (12) and custom python scripts demultiplex and trim the reads based on the inner PCR barcode (1BC) or inner PCR barcode combination (2BC). Lastly, the curated reads for each individual sample were used as an input for Emu (13) (v. 3.4.5) and the default 16S rRNA database (based on rrnDB v.5.6 and NCBI 16S RefSeq from September 2020), for high-confidence species-level annotation. As an output, one receives the relative composition of each input sample depending on the similarity to the full-length 16S sequence. The described Nextflow pipeline is illustrated in **Figure 1**. The Nextflow pipeline was built using singularity containers from https://depot.galaxyproject.org/singularity/. The methodology to process stool sample reads was identical to the processing of bacterial isolate reads except the following differences: The demultiplexed POD5 files were base-called using Dorado (v. 0.5.0; base-caller model “dna_r10.4.1_e8.2_400bps_sup@v4.3.0”; simplex only).

**Figure 1:**
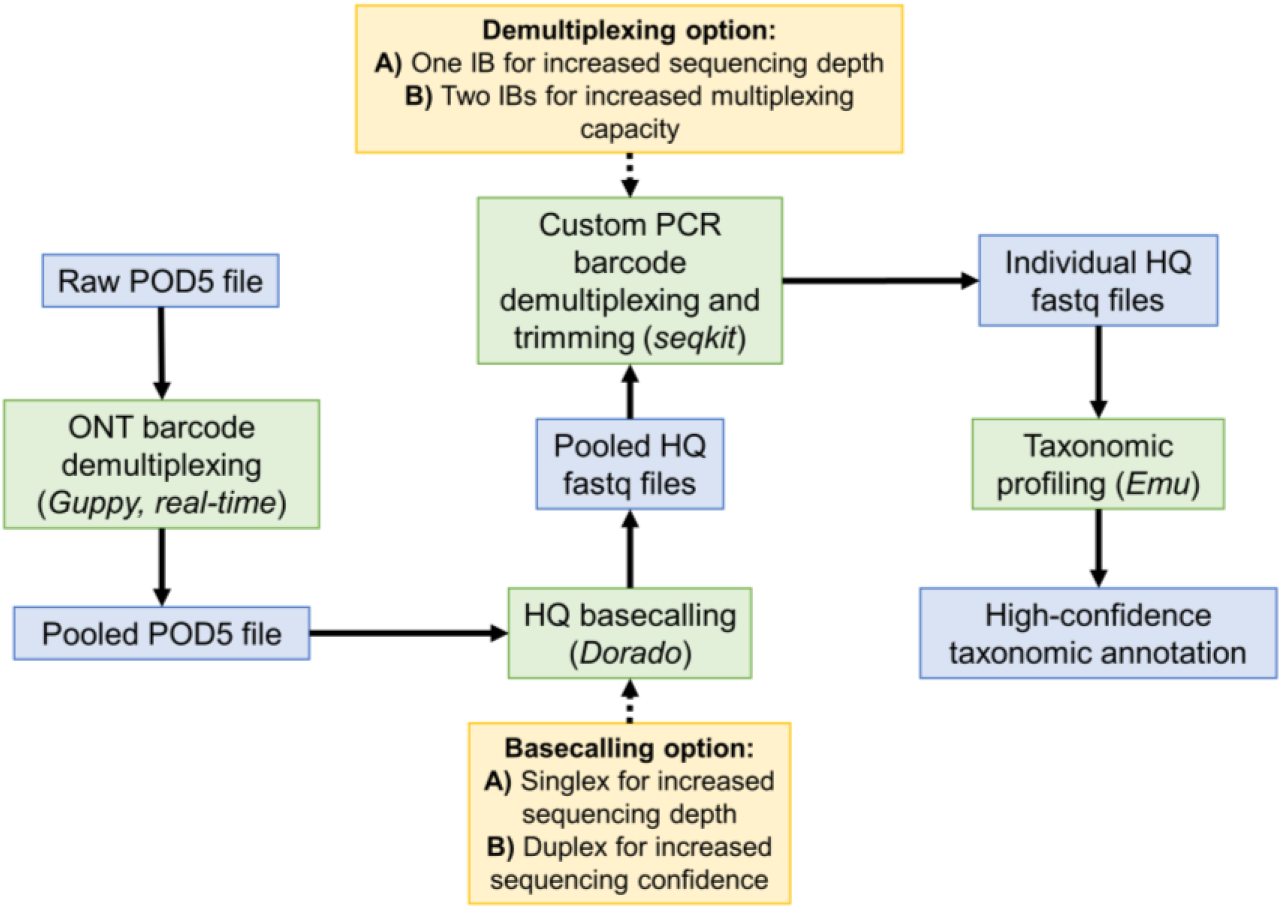
Schematic workflow of the used analysis pipeline. POD5 files were demultiplexed in real-time using Guppy and base-called via Dorado (simplex or duplex). A combination of seqkit and custom python scripts were used to demultiplex based on the inner PCR barcodes (IB) to yield individual high-quality (HQ) fastq files. High confidence-taxonomic annotation was performed using Emu. Obtained files are marked in blue, processing steps in green and processing options in yellow.

### Phylogenetic analysis

To generate per sample consensus sequences for the phylogenetic tree we utilized vsearch (14) (v. 2.23.0) and seqkit (12) (v. 2.5.0) to generate read bins and centroids. Only bins yielding > 10 reads were considered. Subsequently, ONT medaka (v. 1.7.2) was used to generate consensus sequences for each bin and centroid. We used Blast+ (15) (v. 2.13.0) to identify all consensus sequences and select the corresponding sequences for tree building in ClustalW2 (16) (v. 2.1). Bootstrapping (n = 1000) was performed in ClustalW2. The phylogenetic tree was visualized using MEGA-X (v. 10.0.5) and iTOL v5 (17).

### Illumina sequencing and taxonomic profiling of microbial communities

For Illumina sequencing, isolated DNA from stool samples was processed as described in Schneeberger et al. (18). Briefly, samples were sequenced on a Novaseq platform to a depth of >5M paired-end reads per sample in 2x150bp PE mode. Kneadata 0.12.0 was used to filter out any remaining human reads. Metaphlan4 (v. 4.0.4) (19), in combination with the CHOCOPhlAn (vJan21_CHOCOPhlAnSGB_202103) database and Bowtie2 v. 2.4.5 (20), was used to perform taxonomic profiling of the Illumina datasets. The resulting individual profiles were merged using the merge_metaphlan_tables.py script provided with the software.

### Cross-validation via MALDI-TOF MS

To prepare the samples for MALDI-TOF spectra measurement, Columbia agar plates with 5% sheep blood were streaked with the corresponding bacterial stocks. Colonies were left to grow in a CO2 incubator at 37°C and single colonies loaded onto a MALDI-TOF steel target plate using a sterile toothpick. Each colony on the plate was treated with 1μl of 25% formic acid, left to dry and then treated with 1μl of α-Cyano-4-hydroxycinnamic matrix. After sample preparation, the plate was loaded into a microflex® LRF MALDI-TOF mass spectrometer (Bruker, USA) for spectra measurement and analysis (MBT Compass reference library, version 2022). Only samples with ONT annotations > 90% purity (meaning 90% or more reads need to correspond to the same species in the Emu output) were considered and compared to MALDI-TOF MS results. To quantify the strength of the correlation between ONT and MALDI-TOF MS based species annotation, we utilized R (v. 3.4.1) and the R packages vcd (v. 1.4), entropy (v. 1.3.1) and ineq (v. 0.2) to calculate Cramér’s V and Theil’s U. The positive predictive value (PPV) was calculated by dividing the number of true positives (i.e. samples that were identified correctly via MALDI-TOF and our workflow) by the number of true positives plus false positives (i.e. samples that were differently identified by either workflow). As it was not possible to calculate the number of false negatives or true negatives and therefore sensitivity and specificity of our workflow, a PCR run of growth medium negative controls incubated and extracted under the same conditions as the bacterial inoculates was performed. Additionally, a heatmap using R (v. 3.4.1) with the R package pheatmap (v. 1.0.12) visualizing the average percent matches of the ONT reads for each species was generated.

### Processing of microbiome profiles

First, the processed reads of sequencing triplicates were pooled prior to annotation via Emu, resulting in one taxonomic profile for each sample. Secondly, due to the different databases used by the Emu and Metaphlan profilers, we matched the taxonomy of the 16S-based profiles (derived from both 16S primer sets) using the species identified by Illumina as the reference. Briefly, we used a set of criteria to systematically match non-matching species to create a unified abundance file containing all samples and conduct downstream analyses (e.g. analyses of beta diversity). The criteria and their corresponding results are summarized in **Supplementary** Figure 1. Alpha diversity, including the number of taxa, the Shannon diversity, and the Berger-Parker dominance index was calculated for each sample using PAST v. 4.13 (21) and correlated using Spearman correlation available in the XL-STAT software v. 2023.2.1414 (Lumivero 2024, XLSTAT statistical and data analysis solution. https://www.xlstat.com/en). The Bray-Curtis index was used to measure beta diversity and was calculated using the Vegan package v. 2.6-4 (22). The coordinates for the NMDS plot were computed using the metaMDS function from the Vegan package. All figures were generated using OriginPro v. 2024-SR1 (OriginLab Corporation, Northampton, MA, USA).

### Data availability

The Nextflow pipeline is available under: https://github.com/STPHxBioinformatics/ndp. The raw ONT sequencing data (16S rRNA amplicon sequencing) generated in this study have been deposited in the NCBI Short Read Archive under the accession PRJNA1082268.

## Results

### Sequencing performance for bacterial isolates

Overall, we processed 144 bacterial isolates using our ONT-based workflow with the aim to compare its performance for species-level annotation to the current gold-standard technique; MALDI-TOF MS. As shown in **Supplementary Table 7**, after processing of the raw sequencing reads, we obtained average Phred scores of 20+ or 30+ for simplex or duplex data, respectively. For all simplex workflow options (S1BC, S2BC), over 95% samples passed the tentative threshold of 10 reads. Duplex workflows achieved an average of 93% passed samples (D1BC) and 89% passed samples (D2BC), respectively. On average, single barcoding resulted in more reads per sample, compared to double barcoding (S1BC: 2446 and D1BC: 183; S2BC: 2004 and D2BC: 129). Consequently, single barcoding reached higher efficiency values compared to double barcoding (S1BC: 49% and D1BC: 41%; S2BC: 41% and D2BC: 21%). PCR efficiency for each sample pool (sharing the same native ONT barcode) was calculated as the sum of reads containing both a single/double PCR barcode and the native ONT barcode, divided by the total reads for the corresponding native ONT barcode.

### Cross-validation of S2BC data via MALDI-TOF MS

As our workflow will yield more simplex data, which is crucial considering up scalability and multiplexing capabilities, we explored the potential of S2BC reads for bacterial identification. For 47/144 samples, we therefore compared the identification results of our nanopore-based workflow with identification using a MALDI-TOF MS platform. We set a purity cut-off of 90% for each sample - meaning at least 90% of all S2BC reads identified via EMU had to be assigned to the same species. 7/47 samples did not meet this requirement, as less than 90% of annotated reads corresponded to the same species. Identification results from both techniques matched for 36/40 samples. Hence, we obtained a positive predicting value of 0.90 using our nanopore-based workflow and we found strong correlation between both techniques (Cramér’s V = 0.857 and Theil’s U = 0.316). To account for the absence of true negatives in our dataset - and therefore calculation of specificity – a 16S amplicon PCR using growth medium as a negative control was performed, which did not yield any PCR product. To visualize the concordance between the two techniques, we constructed a heatmap (**Figure 2**) showing the average percentage of matching ONT sequencing reads for each tested species. For *E. faecalis, E. hirae, E. coli, L. garviae, and S. anginosus* the average of matching reads lies above 90%.

**Figure 2:**
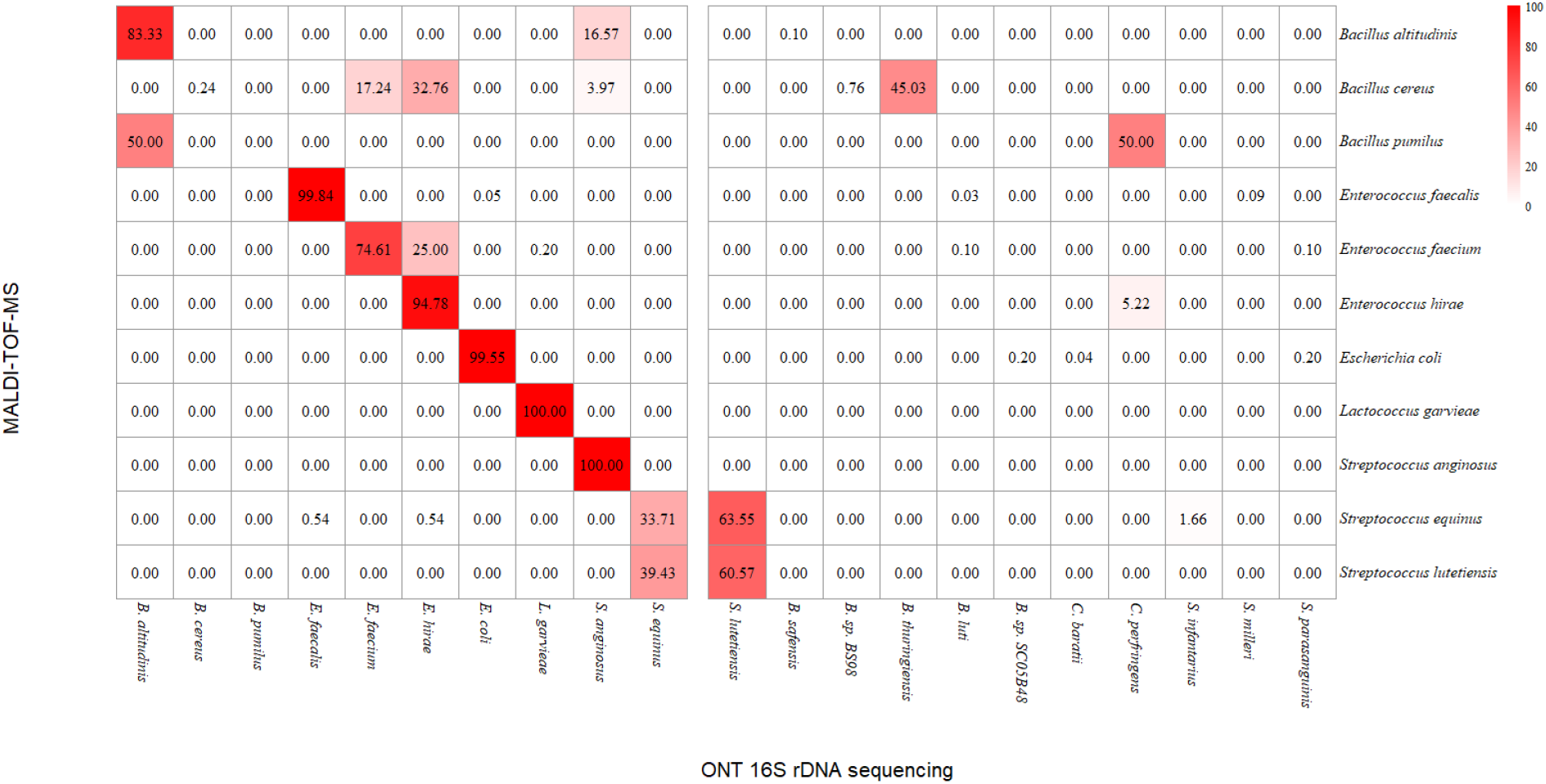
Heatmap visualizing the percentage of ONT 16S rDNA amplicon reads matching the MALDI-TOF MS identification result for each tested species, assuming MALDI-TOF MS as gold standard.

### Identification of bacterial isolates and phylogenetic tree-building using D1BC data

134/144 bacterial isolates (7/144 excluded from MALDI analysis due to contamination and therefore excluded here, 3/144 did not result in any D1BC reads) spanning across 41 species, including closely related and more distant species, were sequenced to assess the taxonomic range of our assay and assess whether D1BC reads were of sufficient quality and quantity for species-level phylogenetic tree-building. We obtained separated clusters on both genus- and species-level, as shown in **Figure 3**.

Furthermore, Emu annotations and blast annotations of the medaka consensus sequences matched in 122/134 cases. Mismatches were comprised of *E. coli* 3, 6-8 (Emu: *E.coli*, blast: *E. fergusonii*), *E.coli* 4-5 (Emu: *E.coli*, blast: *E. marmotae*), *S. oralis* 1 (Emu: *S. oralis*, blast: *S. vulneris*), *S. mitis* 1 (Emu: *S. mitis*, blast: *S. gwangjuense*) and *S. oralis* 3-6 (Emu: *S. oralis*, blast: *S. vulneris*).

**Figure 3:**
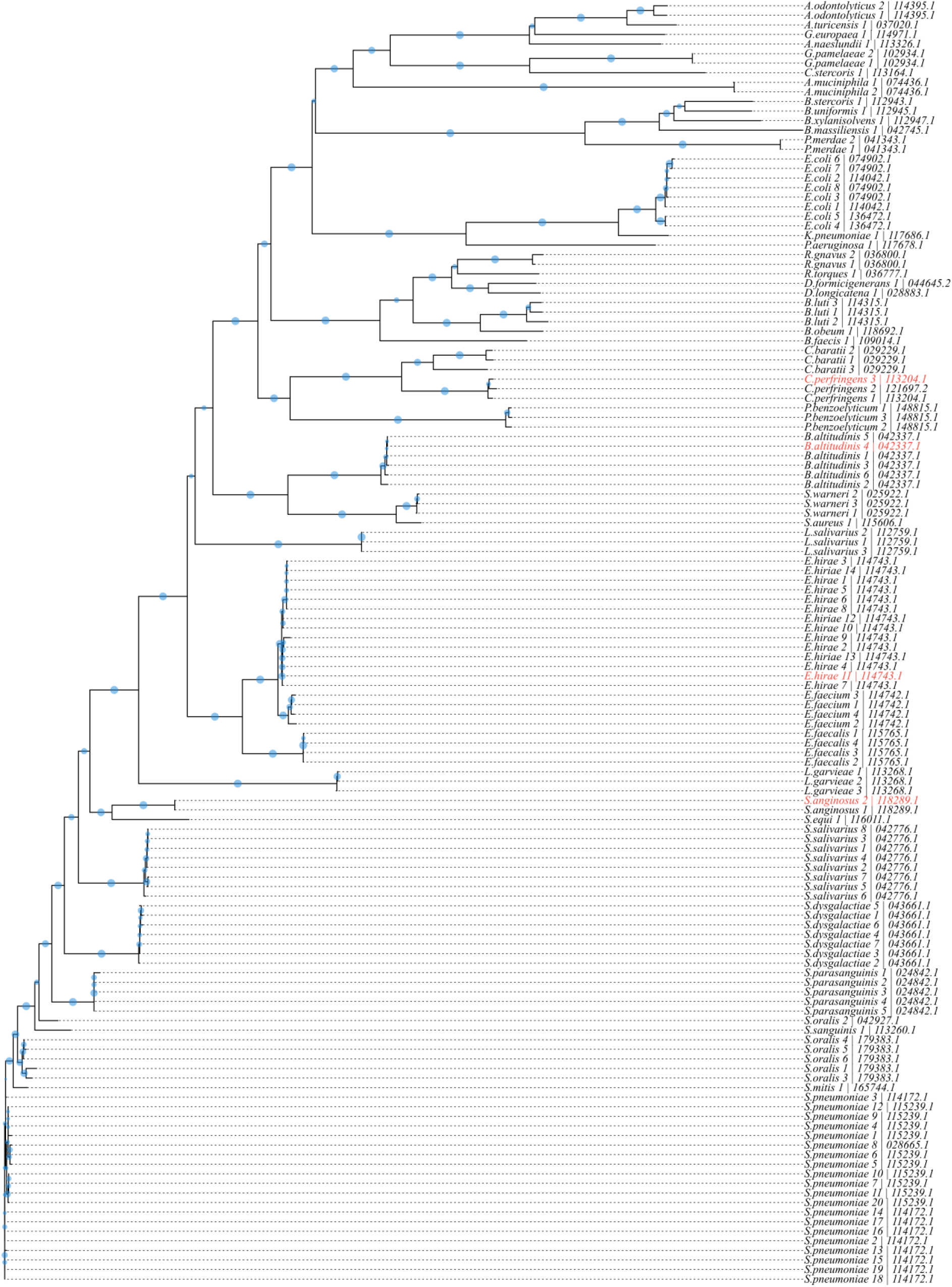
Phylogenetic tree constructed via the multiple sequence alignment of 134 full-length 16S rDNA consensus sequences in ClustalW2. Branch length is based on the distance matrix generated from pairwise alignment scores. Bootstrapping values (n = 1000) are indicated as blue circles on the branches (larger circles correspond to larger bootstrapping values). Isolates marked in red were wrongly identified by our sequencing workflow.

### Performance of full-length 16S sequencing, using two published primer sets, to investigate different features of complex bacterial communities

Using different full-length 16S primer pairs – named A (5) and B (8), we compared the performance of our Nanopore pipeline to investigate species-level microbial community metrics to a standard species-level profiling pipeline based on Illumina sequencing and the widely cited Metaphlan (19) taxonomic profiler. The libraries from 27 stool samples were sequenced in triplicate to mitigate sequencing bias and depth variations arising from flow-cell-specific differences in total sequencing output. An overview of the sequencing run metrics is given in **Supplementary Table 8.** The averaged Phred scores for community sequencing were 30 for A primers and 31 for B primers, respectively (simplex base-calling). Two samples were discarded, one because Illumina library preparation failed and the second because the total read count for 16S sequencing was below 1000. Sequencing depth ranged between 6.4M-18.2M paired-end reads per sample for Illumina, 2081-15920 reads for the A primers, and 1816-22717 reads for the B primers (**Figure 4A**), resulting in 9.73E+08-2.74E+09 base pair (bp) per sample for Illumina, 3E+06-2.31E+07 bp for the A primers, and 2.64E+06-3.28E+07 bp for the B primers. We identified a significant correlation between the number of reads and the number of taxa for the A primer pair (rs = 0.421, *P* = 0.037). However, no such correlation was observed for the other two alpha diversity metrics, namely Shannon diversity (SD) and Berger-Parker (BP) dominance. No correlation was found between sequencing depth and the various alpha diversity measures in the approach based on the B primer pair and Illumina sequencing. Effects of rarefaction for each 16S technique are shown in **Supplementary** Figure 1. When comparing species-level alpha diversity measures, we found a significant correlation between species richness for all comparison pairs, as shown in **Figure 4B**. We also found strong correlation coefficients (r_s_ > 0.6, *P* < 0.001) for each comparison pair when comparing a composite metric of alpha diversity (SD). The dominance index (BP) did not correlate between Illumina and the two 16S primer pairs (r_s_ = 0.361 and 0.337; *P* = 0.077 and *P* = 0.099, respectively) but was strongly correlated between the two 16S-based approaches (r_s_ = 0.839, *P* < 0.001). For inter-samples and inter-techniques comparisons (= beta diversity), the first step was to merge the taxonomy for datasets analyzed with different sequencing techniques and taxonomic profilers (= Illumina versus 16S-based analyses). Since overall diversity was quite low for the ONT-based profiling, we aimed to merge the taxonomy of the 100 most abundant species observed with the A as well as B primers to the taxonomy obtained in the Illumina pipeline. The merging pipeline and the corresponding results are summarized in **Supplementary** Figure 2.

Using the Bray-Curtis dissimilarity measure, we performed a PERMANOVA analysis to compare community-wide composition (**Figure 4C**) and we found that the A-B pair was not significantly different (R^2^ = 0.015 and *P* = 0.687). We found significant differences at the community level between Illumina and pair A (*P* < 0.001) as well as pair B (*P* < 0.001) but low R^2^ values (0.154 and 0.139, respectively) thus indicating that the sequencing technique variable explains only a small proportion of the variability in the taxonomic profiles. Finally, we also compared the Bray-Curtis dissimilarity values of sample pairs across the sequencing techniques and found that pairwise dissimilarity is lower between sample pairs than between unpaired samples for all comparison groups (**Figure 4D**). However, the pairwise dissimilarity is significantly lower between both 16S methods than between Illumina and both 16S sequencing.

**Figure 4:**
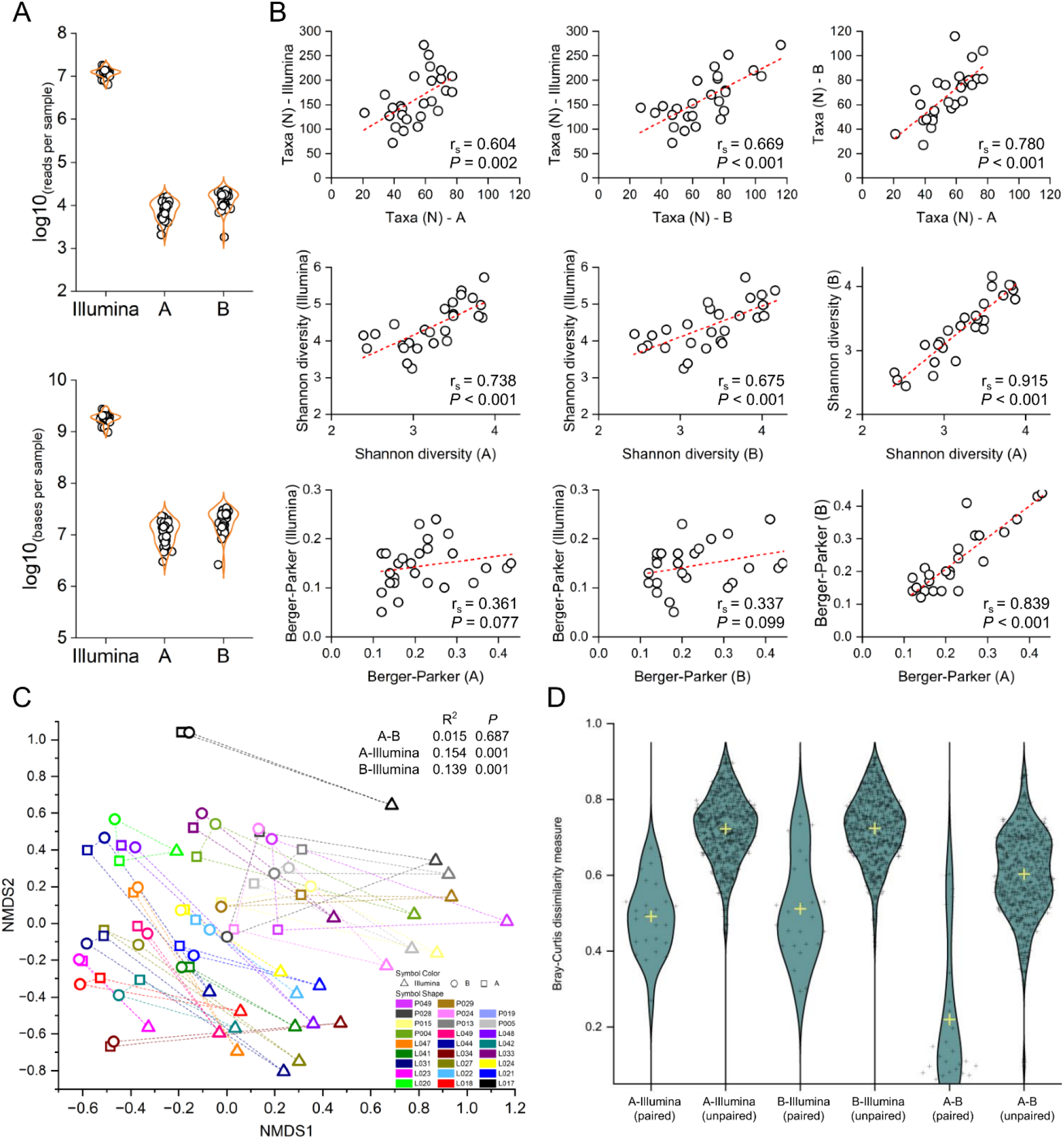
Panel A. Sequencing output per technology in sequenced reads and sequenced base pairs, per sample. **Panel B.** Spearman correlation of alpha diversity metrics between each sequencing technology. **Panel C.** Non-metric multidimensional scaling ordination plot based on Bray-Curtis dissimilarity and results of PERMANOVA analysis. **Panel D.** Bray-Curtis dissimilarity values between paired and unpaired samples and between sequencing technologies.

We then set to provide an in-depth comparison of species-level relative abundances between each analytical strategy. Using the tables with unified taxonomies, we compared the correlation coefficients of each species between the three sequencing protocols (**Figure 5**, left panel). We observed species- and technique-specific variability in the spearman correlation coefficient. Overall, the average Spearman correlation values were similar between Illumina-A and A-B (average(rs) = 0.600 and 0.619, respectively) and slightly lower in the Illumina-B comparison (average(rs) = 0.533), indicating a better agreement in terms of species relative abundances between the Illumina-based and the 16S/A-based profiling. Interestingly, however, there were some differences in terms of presence/absence of different species between the two 16S-based profiling techniques (**Figure 5**, middle panel). For instance, the A primer pair did not pick-up any signal from the *Bifidobacterium* genus, while the B primers were able to detect 4 different *Bifidobacterium* species. Finally, we also showed the fold change for each species between each technique (**Figure 5**, right panel).

**Figure 5:**
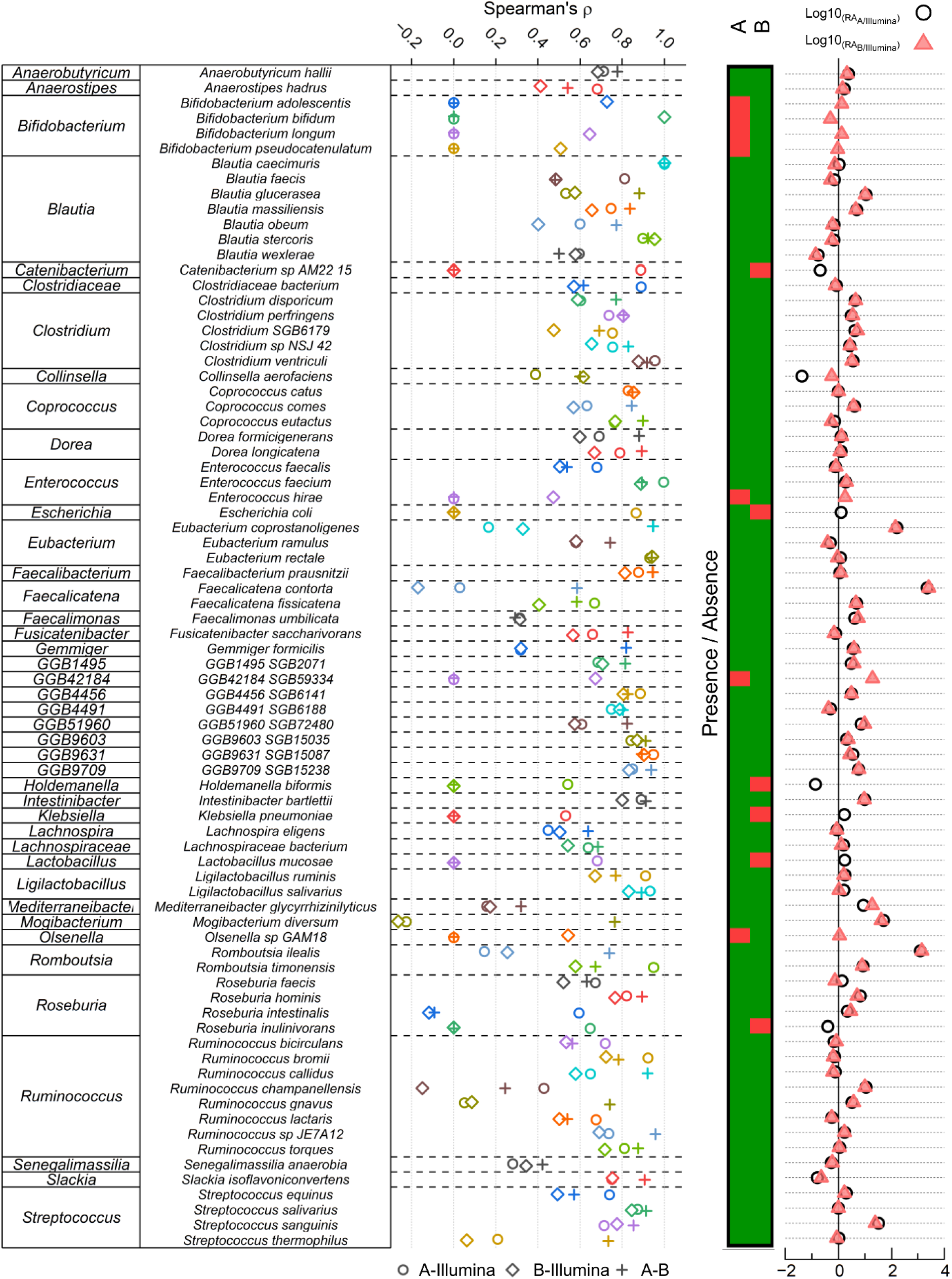
Species-level comparison between sequencing approaches. The left panel indicates the Spearman correlation values of each species between the different sequencing approaches. The middle panel indicates the presence/absence of each species for 16S-based ONT sequencing approaches. Red indicates that the species was not detected, green that it was. All species presented here were detected using Illumina sequencing. The right panel shows the normalized relative abundance fold-change between Illumina and both 16S-based ONT approaches.

### Cost-effectiveness

We estimated current per sample costs for library preparation and sequencing of 80 samples using the ONT Native barcoding kit (SQK-NBD114.96) to approximately 10 USD (**Supplementary Table 9**). By pooling 8 PCR barcoded samples within each ONT barcode, we essentially reduced the sample size to 10. Saved reagent costs result in approximate per-sample costs of 2 USD. In contrast, when characterizing the microbiome of 24 stool samples on a Flongle Flow Cell, we estimate sample cost of roughly 4 USD, compared to 12 USD using the unpooled approach.

## Discussion

Our study presents a novel and flexible approach for sequencing full-length 16S rDNA amplicons using the ONT sequencing platform and the latest v14 kit chemistry. This adaptable workflow yielded promising results across eight sequencing runs, demonstrating the feasibility of the workflow in bacterial identification (2 runs) and microbiome characterization (6 runs) – with a contained cost of around 2 USD per sample for single isolate identification and 4 USD for microbiome characterization.

A major strength of our workflow is its flexibility: If desired, one could adapt the provided code for the use of Emu with other databases such as SILVA (23) or SPINGO (24). Depending on the research question at hand, multiplexing capability, sequencing quality, and read depth can be tuned specifically via the given base calling and barcoding options (S1BC, S2BC, D1BC and D2BC). To each option there are noteworthy consequences: Simplex base-calling will greatly increase read depth, but at the cost of sequencing quality, which appears to be around Q20 (25, 26). While we can expect high quality reads from duplex base-calling, related workflows will result in less reads per sample. Multiplexing more samples via 2 barcodes is possible, at the cost of reduced read depth. Adaptations, such as switching to a SpotOn or PromethION Flow Cells, can enhance throughput in scenarios requiring higher read depth.

While our ONT-results generally align with bacterial species identification data obtained through MALDI-TOF MS, a few divergences were observed, such as misidentification of *B. altitudinis, B. pumilus,* and *E. faecium*. Taxonomically distant misidentifications, such as *B. altitudinis* that was misidentified as *S. lutetiensis*, could be attributed to glycerol stock contamination introduced between MALDI-TOF MS measurements and the sequencing experiment. Interestingly, two misidentifications still shared the correct genus (MALDI: *B. pumilus,* ONT: *B. altitudinis*; MALDI: *E. faecium,* ONT: *E. hirae*). As our workflow was able to identify and distinguish multiple samples belonging to these species, we argue that sequencing quality of these particular samples caused a misidentification.

When comparing full-length 16S sequencing to Illumina sequencing for species-level characterization of microbial communities, we found that while within-sample metrics remained consistent (e.g., Shannon diversity and Berger-Parker index), Illumina-based sequencing detected significantly greater taxonomic richness compared to its 16S-based counterparts. Based on the associations between alpha diversity metrics and sequencing depth found in this dataset, increased per-sample sequencing depth is likely to unveil more low-abundance species, thereby enhancing richness metrics, but at an extra cost linked to decreased multiplexing capacity. Given that many studies and microbiome-based diagnostics prioritize composite metrics (e.g., SD) (27–29), both 16S-based approaches tested in our workflow offer a suitable and cost-effective alternative for characterizing alpha diversity metrics. In terms of inter-sample diversity, and in agreement with previously published studies (30), we observed disparities in taxonomic profiling among the various sequencing technologies (31, 32) and bioinformatics pipelines (33, 34), but the magnitude of these differences remained low (15.4% and 13.9% variability for primers A and B, respectively). While the reasons for differences observed between 16S-based and shotgun-based profiling have been described elsewhere (30), a significant proportion of this variability can be explained with the inability to perfectly match the taxonomies derived from the different analytical pipelines used in this study. Indeed, using our matching criteria, we were only able to match ∼75% of species among the 100 most abundant species observed with 16S-based approaches. Hence, the differences observed in this study are likely over-estimated and using analytical pipelines with unified taxonomies – which are not currently available for both data types – would result in further improved level of agreement between sequencing techniques. We also noted that the two 16S primer pairs yielded very similar results, in accordance with Matsuo et. al (8). It is noteworthy that although the overall composition differed between Illumina and the 16S-based methods, the relative abundances of the most prevalent taxa remained consistent. Therefore, if cost and portability are key factors, and low-abundance taxa are less relevant in specific applications, our proposed approach could be utilized for rapid and cost-effective species-level profiling of complex microbial communities.

At the individual species level, we found good correlation values between Illumina sequencing and the two 16S-based techniques. On average, the A primer showed a better correlation with Illumina in terms of species-level correlation than the B set. However, in terms of presence/absence, the B primer pair picked-up a richer signal, closer to that of Illumina than the A set. For instance, we did not observe any *Bifidobacterium*-related signal in the A dataset but found four *Bifidobacterium* species when using the more recent B primer set, agreeing with the results described by Matsuo et. al (8). Additionally, we observed species-specific disparities in several genera. An example of such is the *Faecalicatena* and *Romboutsia* genera, each represented by two species (*F. contorta* and *F. fissicatena*, and *R. ilealis* and *R. timonensis*). In each case, the normalized fold-change is high (log10(FC) > 3) for one species but low (log10(FC) < 1) for the other species. Hence, while both A and B based approaches are overall comparable in terms of species-level detection, it is essential to carefully weigh these factors and consider their suitability for specific applications.

Our study has several important limitations. Firstly, the selection of bacterial isolates sequenced throughout the study appears to be taxonomically limited and overshadowed by *Enterococcus* and *Streptococcus* species. Therefore, effective detection of untested species might not be guaranteed. However, the agreement of species-level annotations with MALDI-TOF MS data and the robust performance of our workflow in characterization of more complex bacterial communities demonstrated that a vast number of different taxa is in fact detected and correctly annotated. Secondly, we acknowledge that there are substantial differences in sequencing quality of simplex data base-called by Dorado v.0.2.4 (bacterial isolate data; Phred score ∼ 20) versus Dorado v.0.5.0 (stool sample data; Phred score ∼ 30). Despite this difference, simplex data was of adequate quality to annotate bacterial isolate reads with high confidence.

To conclude, our workflow allows for flexible full-length 16S sequencing using recent v14 ONT chemistry with contained sample cost. We demonstrated its robust performance regarding annotation of bacterial isolates (compared to MALDI-TOF MS) and complex bacterial communities (compared to Illumina shotgun sequencing). The provided Nextflow pipeline simplifies data analysis, is modifiable and expandable, while by applying different PCR primers and databases, our protocol can be tailored to diverse research needs. We conclude that S1BC is a suitable option for applications where maximal read depth is needed (e.g. microbiome characterization) and S2BC and D1BC are targeted at large-scale and multiplexed bacterial identification scenarios – leaving the choice between a) more read depth or more multiplexing (S2BC) or b) higher quality reads (D1BC). While modifications are essential for high-throughput applications, the core strengths of our approach lie in its adaptability, flexibility and potential for expansion -hopefully making it a versatile tool for the research community.

## Acknowledgements

We are thankful to Dr. Peter M. Keller and Dr. Carlo Casanova (Institute for Infectious Diseases, University of Berne, Berne, Switzerland), as well as Prof. Dr. Pascale Vonaesch and Dr. Julian Garneau (Department of Fundamental Microbiology, University of Lausanne, Lausanne, Switzerland) who provided several bacterial isolates used in this study. We are thankful to Nurudeen Rahman (Swiss Tropical and Public Health Institute, Allschwil, Switzerland), who contributed to this project with valuable inputs during group discussions. Sincere thanks further go out to Matthias Wurm (Johannes Kepler University, Linz, Austria), who assisted in the development of the analysis pipeline. All steps of the computational analysis were performed using the scientific computing centre at the University of Basel (sciCORE; http://scicore.unibas.ch/). We are grateful to the European Research Council (No. 101019223) for financial support.

## Abbreviations

BHI: Brain Heart Infusion broth
D1BC: duplex with single PCR barcode pair
D2BC: duplex with double PCR barcode pairs
DNA: Deoxyribonucleic acid
HQ: high-quality
IB: Inner PCR barcode
MALDI-TOF: MS Matrix-assisted laser desorption/ionisation, time-of-flight mass spectrometry
ONT: Oxford Nanopore Technologies
PCR: Polymerase chain reaction
RNA: Ribonucleic acid
S1BC: simplex with single PCR barcode pair
S2BC: duplex with single PCR barcode pairs

