## Supplementary Material (PDF) for "A novel barcoded nanopore sequencing workflow of high-quality, full-length bacterial 16S amplicons for taxonomic annotation of bacterial isolates and complex microbial communities"

### Supplementary Figures and Tables

**Supplementary Table 1**: Bacterial isolates used in the study (n = 144).

| Identity | Number of isolates | Origin |
|---|---|---|
| *Actinomyces naeslundii* | 1 | Institute for Infectious Diseases, Berne, Switzerland |
| *Actinomyces odontolyticus* | 2 | Institute for Infectious Diseases, Berne, Switzerland |
| *Actinomyces turicensis* | 1 | Institute for Infectious Diseases, Berne, Switzerland |
| *Akkermansia muciniphila* | 3 | Ordered (https://www.dsmz.de; DSM 22959) |
| *Bacillus altitudinis* | 6 | Isolated from stool at SwissTPH, Allschwil, Switzerland |
| *sp. of Bacillus cereus group* | 2 | Isolated from stool at SwissTPH, Allschwil, Switzerland |
| *Bacillus pumilus* | 2 | Isolated from stool at SwissTPH, Allschwil, Switzerland |
| *Bacteroides massiliensis* | 1 | Ordered (https://www.dsmz.de; DSM 17679) |
| *Bacteroides stercoris* | 1 | Ordered (https://www.dsmz.de; DSM 19555) |
| *Bacteroides uniformis* | 1 | Ordered (https://www.dsmz.de; DSM 6597) |
| *Bacteroides xylanisolvens* | 1 | Ordered (https://www.dsmz.de; DSM 18836) |
| *Blautia faecis* | 1 | Isolated from stool at University of Lausanne, Lausanne, Switzerland |
| *Blautia luti* | 3 | Isolated from stool at University of Lausanne, Lausanne, Switzerland |
| *Blautia obeum* | 1 | Isolated from stool at University of Lausanne, Lausanne, Switzerland |
| *Clostridium baratii* | 3 | Isolated from stool at University of Lausanne, Lausanne, Switzerland (n = 2) Isolated as contaminant at SwissTPH, Allschwil, Switzerland (n = 1) |
| *Clostridium perfringens* | 2 | Isolated from stool at University of Lausanne, Lausanne, Switzerland |
| *Collinsella stercoris* | 1 | Ordered (https://www.dsmz.de; DSM 13279) |
| *Dorea formicigenerans* | 2 | Obtained from University of Lausanne, Lausanne, Switzerland (JCM 31256; n = 1) Ordered (https://www.dsmz.de; DSM 3992, n = 1) |
| *Dorea longicatena* | 1 | Ordered (https://www.dsmz.de; DSM 13814) |
| *Enterococcus faecalis* | 4 | Isolated from stool at SwissTPH, Allschwil, Switzerland |
| *Enterococcus faecium* | 5 | Isolated from stool at SwissTPH, Allschwil, Switzerland |
| *Enterococcus hirae* | 15 | Isolated from stool at SwissTPH, Allschwil, Switzerland |
| *Escherichia coli* | 9 | Isolated from stool at SwissTPH, Allschwil, Switzerland (n = 5) Ordered (https://www.dsmz.de; DSM 30083, n = 1) Isolated from stool at University of Lausanne, Lausanne, Switzerland (n = 3) |
| *Gleimia europaeus* | 1 | Institute for Infectious Diseases, Berne, Switzerland |
| *Gordonibacter pamelaeae* | 2 | Ordered (https://www.dsmz.de; DSM 19378) |
| *sp. of Klebsiella pneumoniae group* | 1 | Ordered (https://www.dsmz.de; DSM 30104) |
| *Lactobacillus salivarius* | 3 | Ordered (https://www.dsmz.de; DSM 20555) |
| *Lactococcus garvieae* | 3 | Isolated from stool at SwissTPH, Allschwil, Switzerland |
| *Parabacteroides merdae* | 2 | Ordered (https://www.dsmz.de; DSM 19495) |
| *Paraclostridium benzoelyticum* | 3 | Isolated from stool at SwissTPH, Allschwil, Switzerland |
| *Pseudomonas aeruginosa* | 1 | Ordered (https://www.dsmz.de; DSM 1128) |
| *Ruminococcus gnavus* | 2 | Ordered (https://www.dsmz.de; DSM 108212) |
| *Ruminococcus torques* | 1 | Isolated from stool at SwissTPH, Allschwil, Switzerland |

| | | |
|---|---|---|
| *sp. of Staphylococcus aureus group* | 1 | Ordered (https://www.dsmz.de; DSM 20231) |
| *Staphylococcus warneri* | 3 | Isolated from stool at SwissTPH, Allschwil, Switzerland |
| *Streptococcus anginosus* | 1 | Isolated from stool at SwissTPH, Allschwil, Switzerland |
| *Streptococcus dysgalactiae* | 7 | Institute for Infectious Diseases, Berne, Switzerland |
| *Streptococcus equi* | 1 | Institute for Infectious Diseases, Berne, Switzerland |
| *Streptococcus equinus* | 2 | Isolated from stool at SwissTPH, Allschwil, Switzerland |
| *Streptococcus lutetiensis* | 1 | Isolated from stool at SwissTPH, Allschwil, Switzerland |
| *Streptococcus mitis* | 1 | Institute for Infectious Diseases, Berne, Switzerland |
| *Streptococcus oralis* | 6 | Institute for Infectious Diseases, Berne, Switzerland |
| *Streptococcus parasanguinis* | 5 | Ordered (https://www.dsmz.de; DSM 6778) |
| *Streptococcus pneumoniae* | 20 | Institute for Infectious Diseases, Berne, Switzerland |
| *Streptococcus salivarius* | 8 | Ordered (https://www.dsmz.de; DSM 20067) |
| *Streptococcus sanguinis* | 1 | Isolated from stool at University of Lausanne, Lausanne, Switzerland |

**Supplementary Table 2:** List of primers (A) used for the amplification of bacterial 16S rDNA in bacterial isolate samples and stool samples that were adapted from Urban et al. (7). Indicated in bold are the 12 bp long barcodes used for demultiplexing, followed by either a 27F or 1492R binding sequence.

| Primer | Primer Sequence |
|---|---|
| P01-FWD | **GAGCCCGTTCCG**AGAGTTTGATCMTGGCTCAG |
| P02-FWD | **TGGCACCGATTA**AGAGTTTGATCMTGGCTCAG |
| P03-FWD | **GACATACAATGA**AGAGTTTGATCMTGGCTCAG |
| P04-FWD | **ATGGTCTACTAC**AGAGTTTGATCMTGGCTCAG |
| P05-FWD | **CCACTTGGATAG**AGAGTTTGATCMTGGCTCAG |
| P06-FWD | **CGATTATGGCAC**AGAGTTTGATCMTGGCTCAG |
| P07-FWD | **CTTACGAGGCAT**AGAGTTTGATCMTGGCTCAG |
| P08-FWD | **GTCCACCCTGGG**AGAGTTTGATCMTGGCTCAG |
| P01-REV | **GAGCCCGTTCCG**CGGTTACCTTGTTACGACTT |
| P02-REV | **TGGCACCGATTA**CGGTTACCTTGTTACGACTT |
| P03-REV | **GACATACAATGA**CGGTTACCTTGTTACGACTT |
| P04-REV | **ATGGTCTACTAC**CGGTTACCTTGTTACGACTT |
| P05-REV | **CCACTTGGATAG**CGGTTACCTTGTTACGACTT |
| P06-REV | **CGATTATGGCAC**CGGTTACCTTGTTACGACTT |
| P07-REV | **CTTACGAGGCAT**CGGTTACCTTGTTACGACTT |
| P08-REV | **GTCCACCCTGGG**CGGTTACCTTGTTACGACTT |

**Supplementary Table 3:** List of primers (B) used for the amplification of bacterial 16S rDNA in stool samples that were adapted from Matsuo et al. (10) and were used for microbiome characterization. Indicated in bold are the 12 bp long barcodes used for demultiplexing, followed by either a 27F or 1492R binding sequence containing multiple degenerate bases (R, Y, M).

| Primer | Primer Sequence |
|---|---|
| P01-FWD | **GAGCCCGTTCCG**AGRGTTYGATYMTGGCTCAG |
| P02-FWD | **TGGCACCGATTA**AGRGTTYGATYMTGGCTCAG |
| P03-FWD | **GACATACAATGA**AGRGTTYGATYMTGGCTCAG |
| P04-FWD | **ATGGTCTACTAC**AGRGTTYGATYMTGGCTCAG |
| P05-FWD | **CCACTTGGATAG**AGRGTTYGATYMTGGCTCAG |
| P06-FWD | **CGATTATGGCAC**AGRGTTYGATYMTGGCTCAG |
| P07-FWD | **CTTACGAGGCAT**AGRGTTYGATYMTGGCTCAG |
| P08-FWD | **GTCCACCCTGGG**AGRGTTYGATYMTGGCTCAG |
| P01-REV | **GAGCCCGTTCCG**CGGYTACCTTGTTACGACTT |
| P02-REV | **TGGCACCGATTA**CGGYTACCTTGTTACGACTT |
| P03-REV | **GACATACAATGA**CGGYTACCTTGTTACGACTT |
| P04-REV | **ATGGTCTACTAC**CGGYTACCTTGTTACGACTT |
| P05-REV | **CCACTTGGATAG**CGGYTACCTTGTTACGACTT |
| P06-REV | **CGATTATGGCAC**CGGYTACCTTGTTACGACTT |
| P07-REV | **CTTACGAGGCAT**CGGYTACCTTGTTACGACTT |
| P08-REV | **GTCCACCCTGGG**CGGYTACCTTGTTACGACTT |

**Supplementary Table 4:** Reagents and volumes used per well in the 16S rDNA PCR reaction. Primers and DNA template were diluted in nuclease-free water.

| Reagent | Volume per reaction [µl] |
|---|---|
| Nuclease-free water | 8.5 |
| NEB LongAmp Hot Start *Taq* 2x Master Mix | 12.5 |
| 16S forward primer (10µM) | 1 |
| 16S reverse primer (10µM) | 1 |
| DNA template (1:10 diluted) | 2 |
| Reaction volume | 25 |

**Supplementary Table 5:** PCR setup conditions used for the 16S amplicon barcoding reaction.

| Cycling conditions | | |
|---|---|---|
| **Temperature** | **Time** | |
| 94°C | 30s | |
| 94°C | 30s | 30-35 x |
| 65°C | 15s | |
| 65°C | 50s | |
| 65°C | 10min | |

**Supplementary Table 6:** Chosen settings for performed sequencing runs using a Flongle Cell on a MinION Mk1C. Runtime was entirely dependent on the Flongle Flow cell and number of total reads reaching a plateau phase.

| | |
|---|---|
| **Runtime** | 10 - 22h |
| **Minimum length** | 200bp |
| **Minimum Q-score** | 9 |
| **Base-calling** | Fast Base-calling |
| **Barcoding** | On (default) |
| **File: Raw-reads** | POD5 |
| **File: Base called** | FastQ |

**Supplementary Table 7:** Overview of the two performed bacterial isolate sequencing runs for the four base-calling and barcoding options (S = simplex; D = duplex; S1BC = simplex, 1 PCR barcode; S2BC = simplex, 2 PCR barcodes; D1BC = duplex, 1 PCR barcode; D2BC = duplex, 2 PCR barcodes). Samples with more than 10 reads were classified as "Passed samples".

| **Run metrics** | | | **average** |
|---|---|---|---|
| Primer set | A | A | |
| Number of pooled samples (n) | 75 | 75 | |
| Number of initial flongle pores | 78 | 75 | |
| Number of sequenced nucleotides [Mb] | 1270 | 634 | 952 |
| Number of S / D reads (before filtering) | 537464 / 67985 | 268509 / 39607 | 402987 / 53796 |
| Number of S / D reads (after filtering) | 482835 / 64492 | 248294 / 37834 | 365565 / 51163 |
| Average S / D Phred score (after filtering) | 22 / 30 | 21 / 31 | 22 / 31 |

| **S1BC** | | | **average** |
|---|---|---|---|
| Average reads per sample | 3289 | 1603 | 2446 |
| Average barcoding efficiency [%] | 50 | 48 | 49 |
| Passed samples | 74 | 73 | 73.5 |
| Passed samples [%] | 98.7 | 97.3 | 98.0 |

| **S2BC** | | | **average** |
|---|---|---|---|
| Average reads per sample | 2678 | 1329 | 2004 |
| Average barcoding efficiency [%] | 41 | 40 | 41 |
| Passed samples | 74 | 72 | 73 |
| Passed samples [%] | 98.7 | 96.0 | 97.3 |

| **D1BC** | | | **average** |
|---|---|---|---|
| Average reads per sample | 227 | 139 | 183 |
| Average barcoding efficiency [%] | 28 | 30 | 29 |
| Passed samples | 71 | 68 | 69.5 |
| Passed samples [%] | 94.7 | 90.7 | 92.7 |

| **D2BC** | | | **average** |
|---|---|---|---|
| Average reads per sample | 158 | 100 | 129 |
| Average barcoding efficiency [%] | 20 | 22 | 21 |
| Passed samples | 69 | 64 | 66.5 |
| Passed samples [%] | 92.0 | 85.3 | 88.7 |

**Supplementary Table 8:** Overview of the six performed stool sample sequencing runs using the S1BC base-calling and barcoding option (A = full-length 16S primers adapted from Urban et al. (7); B = full length 16S primers adapted from Matsuo et al. (10); S1BC = simplex, 1 PCR barcode).

| **Run metrics** | | | | | | | **average (A)** | **average (B)** |
|---|---|---|---|---|---|---|---|---|
| Primer set | A | A | A | B | B | B | | |
| Number of pooled samples (n) | 27 | 27 | 27 | 27 | 27 | 27 | | |
| Number of initial flongle pores | 76 | 73 | 69 | 66 | 89 | 103 | 73 | 86 |
| Number of simplex reads (before filtering) | 200819 | 113003 | 265733 | 198244 | 307634 | 296618 | 193185 | 267499 |
| Number of simplex reads (after filtering) | 159607 | 103400 | 164508 | 152833 | 236713 | 195376 | 142505 | 194974 |
| Average simplex Phred score (after filtering) | 31 | 31 | 28 | 30 | 31 | 32 | 30 | 31 |
| Average S1BC reads per sample | 3429 | 2145 | 3229 | 3604 | 5674 | 4623 | 2934 | 4634 |
| Average S1BC barcoding efficiency [%] | 58% | 56% | 53% | 65% | 66% | 65% | 56% | 65% |

**Supplementary Table 9:** Estimated per sample cost for library preparation and sequencing of 24 (e.g. for microbiome characterization) or 80 (e.g. for single isolate annotation) samples using pooled approaches or unpooled approaches.

| **Estimated sequencing costs by approach** | | | **per sample costs** | |
|---|---|---|---|---|
| **Reagent** | **kit costs** | **kit reactions** | **unpooled (n)** | **pooled (n / 8)** |
| Native Barcoding Kit 96 v14 | $ 799.00 | 288 | $ 2.77 | $ 0.35 |
| NEBNext® dA-Tailing Module (E6053L) | $ 650.00 | 384 | $ 1.69 | $ 0.21 |
| Blunt/TA Ligase Master Mix (M0367L) | $ 600.00 | 125 | $ 4.80 | $ 0.60 |
| NEBNext® Quick Ligation Module (E6056L) | $ 1’950.00 | 8000 | $ 0.24 | $ 0.24 |
| cost per reaction | | | $ 9.51 | $ 1.40 |
| cost flongle flow cell | | | $ 65.00 | $ 65.00 |
| reaction cost for n = 80 samples | | | $ 825.86 | $ 177.17 |
| per sample cost for n = 80 samples | | | **$ 10.32** | **$ 2.21** |
| reaction cost for n = 24 samples | | | $ 293.26 | $ 98.65 |
| per sample cost for n = 24 samples | | | **$ 12.22** | **$ 4.11** |

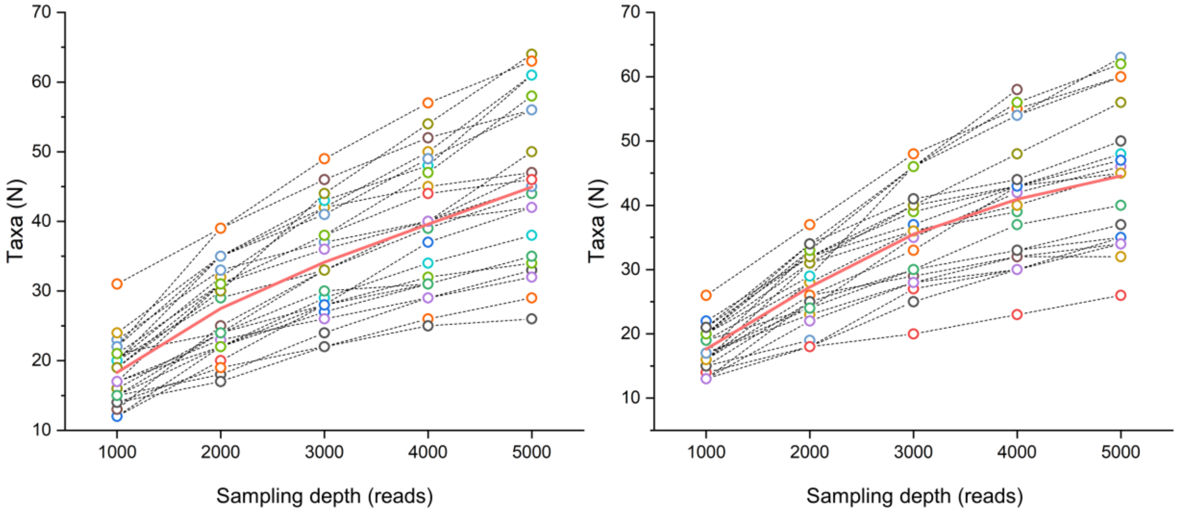


**Supplementary Figure 1:** Rarefaction curves of the 16S-based approaches, using A (left panel) and B (right panel) primer pairs.

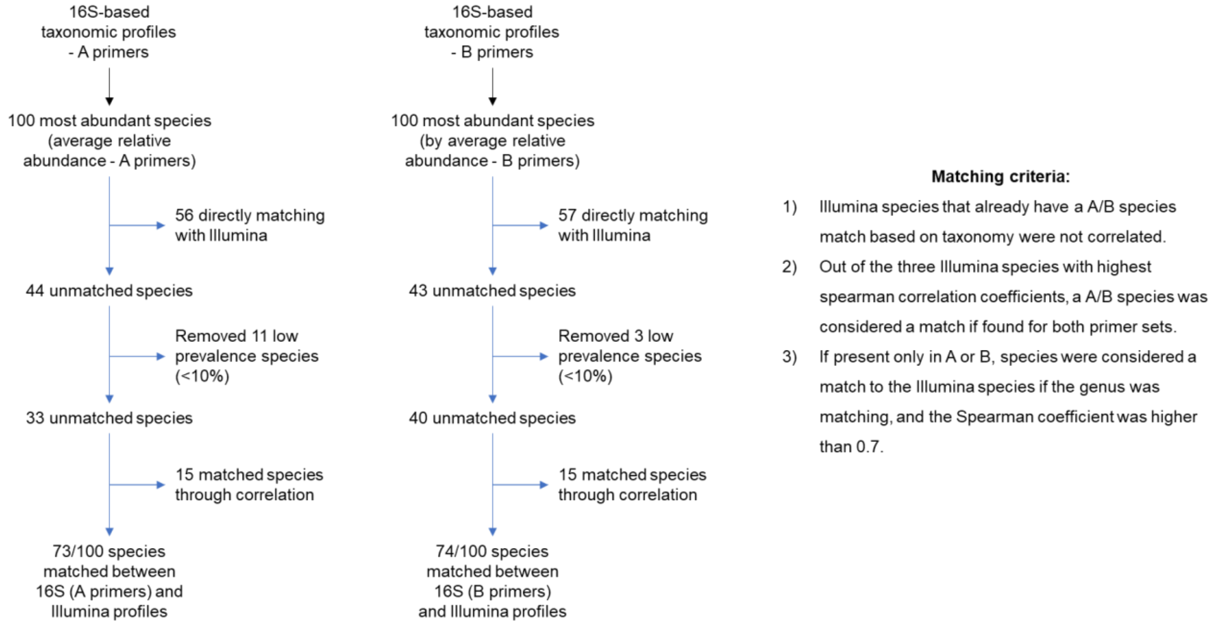


**Supplementary Figure 2:** Proportion of matched species between 16S-based and Illumina-based results and criteria used to merge divergent taxonomies resulting from different taxonomic profilers.
